# NONO as a Sensor of Intracellular Oxidation: Relevance to Neuroblastoma Cell Death

**DOI:** 10.1101/2025.10.30.685543

**Authors:** Sofya S. Pogodaeva, Olga O. Miletina, Nadezhda V. Antipova, Alexander A. Shtil, Oleg A. Kuchur

**Affiliations:** National Research University ITMO, 197101 Saint-Petersburg, Russia; Shemyakin-Ovchinnikov Institute of Bioorganic Chemistry, Russian Academy of Sciences, 117997 Moscow, Russia; Blokhin National Medical Research Center of Oncology, 115522 Moscow, Russia; Higher School of Economics, 194100 Saint-Petersburg, Russia

**Keywords:** core regulatory circuit, MYCN, NONO, auranofin, neuroblastoma

## Abstract

Neuroblastoma, a transcriptionally driven pediatric malignancy, exhibits a remarkable clinical and biological heterogeneity. Two major subtypes, the adrenergic and mesenchymal, are differentially governed by a subset of transcription factors that comprise the core regulatory circuit (CRC). The former subtype is often associated with *MYCN* amplification and is particularly aggressive and therapy-resistant, underscoring the need for novel targets. Here, we identify the multifunctional non-POU domain-containing octamer-binding (NONO) protein as a guardian of individual CRC genes, thereby contributing to survival of neuroblastoma cells with different *MYCN* copy numbers. Intracellular oxidation in response to auranofin, an inhibitor of thioredoxin reductase 1, rapidly down-regulated the amounts of NONO mRNA and protein in *MYCN*-amplified Kelly cell line. Conversely, *NONO* knockdown with RNA interference (siNONO) also triggered intracellular oxidation. These effects were less pronounced in the SK-N-AS cell line carrying a single *MYCN* copy, as well as in non-malignant HS5 fibroblasts. In Kelly cells, siNONO attenuated auranofin-induced activation of CRC genes *HAND2* and *PHOX2B*. In line with preferential effects on NONO abundance, the Kelly cells were more sensitive than single *MYCN* copy counterparts to combinations of a sublethal concentration of auranofin with siNONO. Importantly, *MYCN*-amplified cells demonstrated a significantly suppressed clonogenic survival 14 days after transient exposure to these combinations compared with each agent alone; HS5 fibroblasts were largely spared. Our findings 1) establish NONO as a redox sensor, a non-trivial role for transcriptional proteins, and 2) justify the strategy of therapeutic targeting of *MYCN*-amplified tumors vulnerable to oxidative stress.

**Key points:**

- NONO, a master regulator of the core regulatory circuit (CRC) in *MYCN*-amplified neuroblastoma, is rapidly down-regulated by auranofin-induced intracellular oxidation.
- NONO knockdown synergizes with auranofin in triggering individual CRC gene deregulation and lethal oxidative stress preferentially in *MYCN*-amplified cells.

## Introduction

Neuroblastoma (NB) is a pediatric malignancy of the sympathetic nervous system, arising from embryonic neural crest precursors. It shows extreme clinical heterogeneity: some tumors spontaneously regress, whereas others progress relentlessly despite intensive multimodal therapy [1–3]. About half of patients present with high-risk disease, for which 5-year survival remains below 50 % [4]. Like most childhood tumors, NB has a low somatic mutation burden; instead, widespread epigenetic and transcriptional deregulation are predominant. A hallmark of high-risk NB is amplification of the *MYCN* oncogene, an established marker of the aggressive disease. Other factors such as activating *ALK* mutations or 11q deletions occur in subsets of cases [5–8]. Thus, NB is better characterized by a ‘transcriptional burden’ [9], underscoring the importance of gene-regulatory therapeutic programs.

Genome-wide expression profiling has revealed two major NB cell states termed the adrenergic (ADRN) and mesenchymal (MES) subtypes [10, 11]. The ADRN subtype is typically linked to *MYCN* gene amplification and is defined by a cohort of transcription factors (*PHOX2B, HAND2, GATA3, ISL1, TBX2, ASCL1*, etc.) that regulate each other at the level of gene expression. These factors form a self-reinforcing core regulatory circuit (CRC) [12, 13]. In contrast, MES cells express a different lineage program, e.g., PRRX1, YAP/TAZ and SNAI1, and resemble neural-crest–derived mesenchymal phenotypes. This state tends to be more chemoresistant and invasive, and may expand after treatment pressure [14–16]. Importantly, both states can undergo transition and transdifferentiate in the course of therapy [17, 18]. This transcriptional heterogeneity and plasticity, along with low immunogenicity, complicates the treatment.

Current regimens for high-risk NB include intensive chemotherapy, radiotherapy, anti-GD2 immunotherapy and autologous stem-cell rescue [19–21]. However, the majority of tumors relapse. Over half of children with high-risk NB eventually experience disease recurrence, which significantly reduces survival rates [22, 23]. Conventional therapies do not specifically address the individual molecular ‘portraits’ of NB subtypes; therefore, the tumors can escape standard cytotoxic agents [24–26]. These limitations highlight the need for novel approaches that target the biological peculiarities of NB.

CRC targeting has emerged as an attractive strategy. The genome-wide screens showed that MYCN, HAND2, ISL1, PHOX2B, GATA3 and TBX2 transcription factors are essential for *MYCN*-amplified NB [27, 28]. Disruption of this circuit profoundly impairs NB growth. Combined inhibition of transcriptional proteins BRD4 and CDK7 caused rapid loss of CRC transcription and synergistically suppressed *MYCN*-amplified tumors [29]. These reports as well as other studies emphasized a network of distinct CRC members and auxiliary survival pathways that act in concert to promote NB cell survival (reviewed in [30]).

Among transcriptional mechanisms in NB, the non-POU domain-containing octamer-binding protein (NONO or p54nrb) deserves a special heed. NONO, a multifunctional DNA/RNA-binding protein, has been recently identified as a master regulator of oncogenic transcription in NB [31, 32]. NONO binds to 5′-ends of introns and to long non-coding RNAs (lncRNAs) including *MYCN* transcript, promoting the expression of oncogenes [33]. Through these interactions, NONO ‘guards’ CRC and amplifies MYCN signaling. Clinically, high NONO expression correlates with aggressive NB and poor prognosis [34].

Thus, NONO exemplifies a critical regulator of NB states whose pharmacologic inhibition is desirable. However, NONO has long been considered undruggable since this protein lacks an enzymatic structure; only very recently the small molecular weight compound that disrupts NONO-RNA interaction has been introduced [35–37] (*36, 37 not yet in an issue*). Nevertheless, evidence has been reported in favor of NONO inhibition as a result of metabolic deregulation.

Auranofin is a clinically approved irreversible inhibitor of the antioxidant enzyme thioredoxin reductase 1 (TrxR1) [38]. Consequently, auranofin forces intracellular accumulation of reactive oxygen species (ROS). This effect has been used for induction of oxidative stress in neurogenic tumors [39–41]. Notably, *MYCN*-amplified NB cells rely on robust redox buffers; therefore, these cells may be particularly vulnerable to the shift of redox balance to oxidation [42]. Intriguingly, Hou et al. have shown that auranofin can inactivate NONO in glioblastoma [43]. Although the mechanism of inhibition remained to be elucidated, this pivotal observation set the stage for a detailed investigation of the role of NONO in specific metabolic responses as well as for therapeutic combinations to enhance intracellular oxidation in NB cells.

In this study, we investigated whether intracellular oxidation is mechanistically linked to NONO inactivation, deregulation of individual CRC genes and death of NB cells carrying different numbers of *MYCN* copies. The CRC genes *PHOX2B* and *HAND2* were chosen because the former transcription factor reflects the integrity of ADRN lineage-determining program whereas *HAND2* participates in enhancer-dependent transactivation and MYCN-mediated proliferation [28, 44].

## Materials and Methods

### Chemicals

Reagents were purchased from Dia-M, Russia unless specified otherwise.

### Cell culture

Human NB cell lines with (Kelly and IMR-32) or without (SK-N-AS) *MYCN* amplification were used. These cell lines were purchased from the American Type Culture Collection (Manassas, VA). The human bone marrow-derived fibroblast cell line HS5 (gift of Dr. T. Lebedev, Engelhardt Institute of Molecular Biology, Russian Academy of Sciences, Moscow) was taken as a non-transformed counterpart. The Kelly and HS5 cell lines were propagated in RPMI-1640; IMR-32 cells were grown in Eagle’s minimum essential medium supplemented with 10% FBS and 100 µg/mL gentamicin. The SK-N-AS cells were cultured in Dulbecco’s modified Eagle medium with high glucose and 1% non-essential amino acids. Cells were cultured at 37 °C, 5% CO_2_ in a humidified atmosphere. Cells were routinely tested and found to be free from *Mycoplasma* contamination.

### Compounds and cell treatment

Auranofin (Astellas Pharma, Japan) was dissolved in 10 mM dimethyl sulfoxide and stored at −20^0^C until the experiments. For transient gene knockdown, five siRNAs were used: *NONO, MYCN, HAND2, PHOX2B* (CRC gene-specific) and siRNA to *GFP* (unrelated control). All siRNAs (DNA Synthesis, Russia; Table 1) were transfected with GenJect liposomes 40 (Molecta, Russia): 2 µL siRNA, 125 nM liposomes in 35 mm Petri dish for 24 h according to the manufacturer’s instructions. Gene knockdown was registered 24 h post transfection. Two to three variants of each siRNA sequence were tested to achieve the most potent and reproducible knockdown. In combination experiments, cells were first transfected with the respective siRNAs; 24 h later the medium was removed, and monolayers were washed with PBS. Fresh medium supplemented with auranofin was added, and cells were incubated followed by cytotoxicity assays (see below).

**Table 1.**
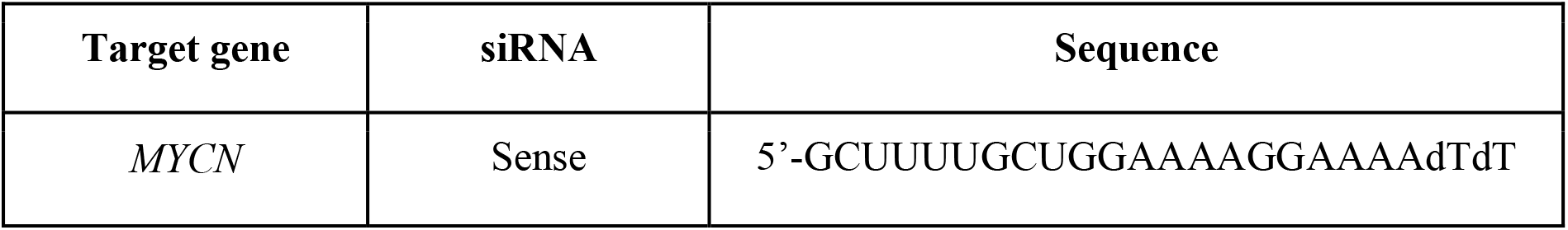
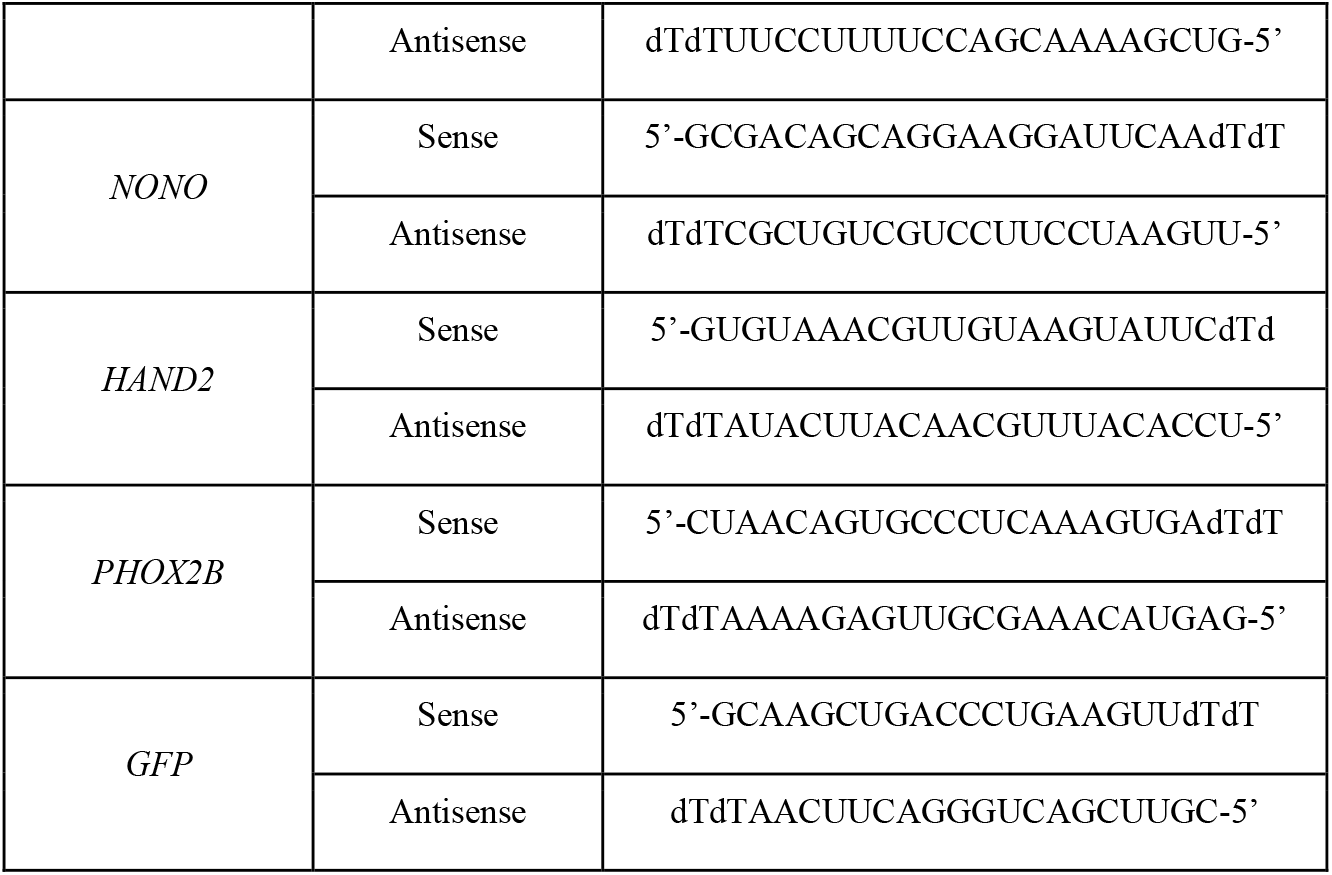
Sequences of siRNAs.

### Intracellular detection of reactive oxygen species

After incubation with auranofin without or with siRNAs, cells were loaded with 5 µM 2′,7′-dichlorodihydrofluorescein diacetate (DCFH_2_-DA), incubated for 60 min at 37 °C, then washed with PBS, harvested by centrifugation (500g, 5 min) and resuspended in 200 µL phosphate buffered saline (PBS). Fluorescence intensity was analyzed on a CytoFlex B2-R2-V0 flow cytometer (Beckman Coulter, Brea, CA) (excitation/emission settings 493/523 nm; FITC channel, 488 nm laser). Hydrogen peroxide (0.5 mM) was used as a control for induction of intracellular oxidation.

### Total RNA extraction and qRT-PCR analysis

Isolation of total RNA from cells was performed with the ExtractRNA buffer (Evrogen, Russia) according to the manufacturer’s protocol. cDNA was synthesized from 2 μg RNA preparation using MMLV reverse transcriptase: 25°C 10 min, 42°C 50 min, 70°C 10 min, 10°C 10 sec. The PCR mixture contained 1 μg cDNA, 5x qPCR SYBR Green I, forward and reverse primers (10 μM each; Table 2), and nuclease-free H2O.

**Table 2.**
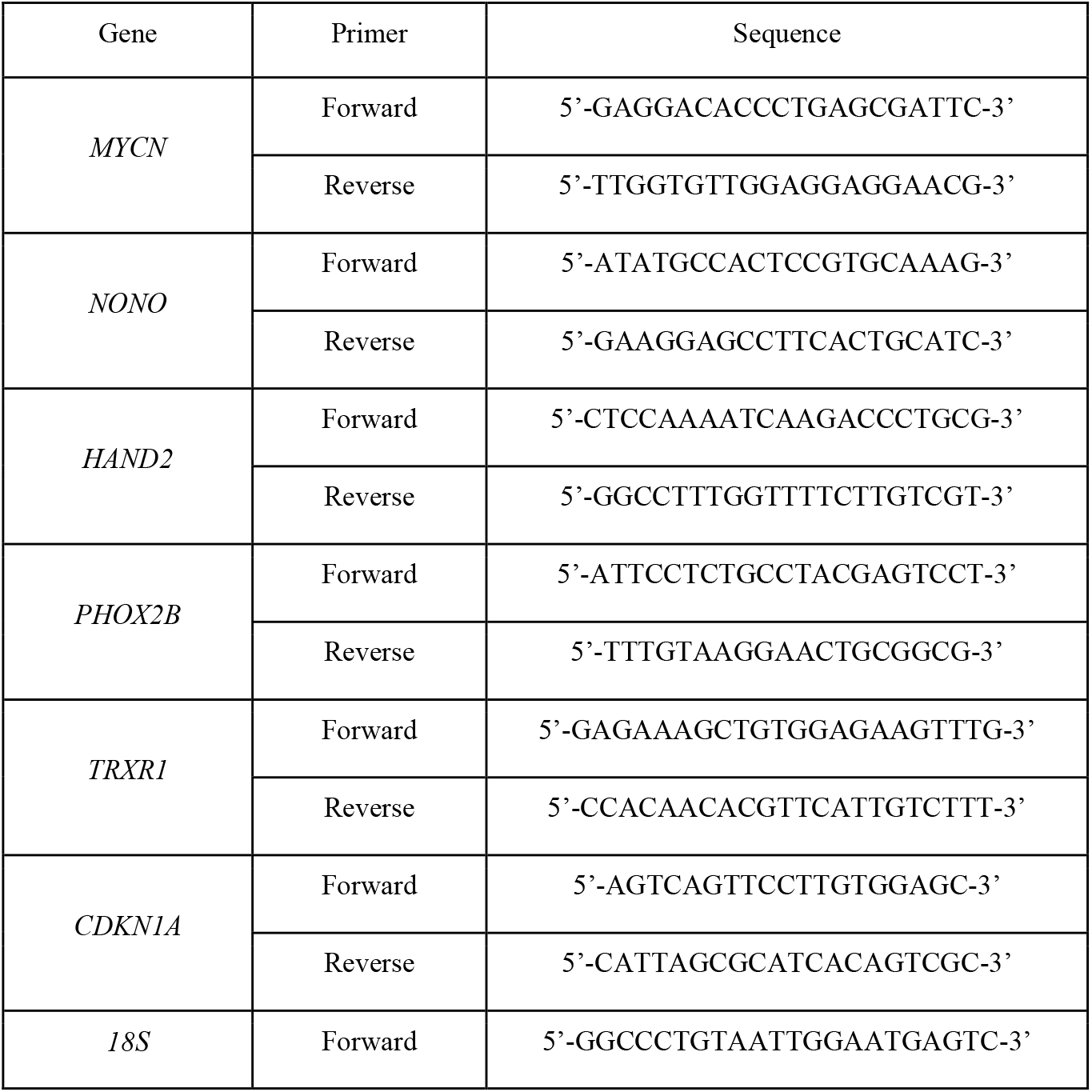
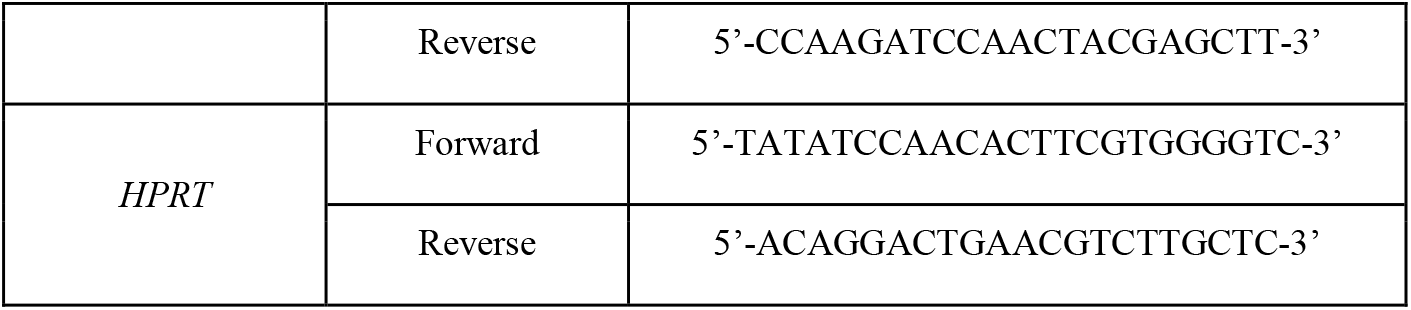
Primers for RT-PCR.

Amplification conditions:

- Stage 1 (1 cycle): 94°C 3 min, 60°C 40 sec, 72°C 40 sec
- Stage 2 (28–30 cycles): 94°C 10s, 60°C 10s, 72°C 20 sec
- Stage 3 (1 cycle): 72°C 3 min
- Stage 4 (storage): 4°C

After the completion of reactions, the relative expression was determined by the ΔCt method, where Ct (threshold cycle) is the cycle at which the fluorescence level reaches a certain value (preselected threshold), and Δ is the change in the expression of the gene of interest relative to the transcript selected for normalization (hypoxanthine-guanine phosphoribosyltransferase (*HPRT*) and *18S RNA* genes; difference < 2 cycle).

### Immunoblotting

After the completion of treatments, cells were lysed in a buffer containing 50 mM Tris-HCl pH 8.0, 150 mM NaCl, 1% NP-40, 0.1% sodium dodecyl sulfate, 200 μM phenylmethylsulfonyl fluoride, protein inhibitor cocktail (leupeptin, aprotinin and pepstatin, 1 μg/ml each) (Sigma-Aldrich, Burlington, MA). Protein concentration was measured by Bradford assay. Lysates (35 µg total protein per lane) were separated by electrophoresis in 10% sodium dodecyl sulfate-polyacrylamide gels and transferred onto a nitrocellulose membrane (BioRad, Hercules, CA) at 250 mA, 1 h at 4 °C. Membranes were blocked with 5% bovine serum albumin for 30 min at room temperature and incubated with primary antibodies (1:1000) overnight at 4 °C. After washing with Tris-buffered saline-Tween-20, membranes were incubated with horseradish peroxidase-conjugated secondary antibodies (1:1000, Cell Signaling Technology, Danvers, MA) for 1 h. Signals were detected with Enhanced Chemolumiscence reagent (Thermo Fisher, Waltham, MA) and captured using a gel documentation system ChemiDoc™ Touch (BioRad, Hercules, CA) with multiple exposure times to ensure linear range of detection. The primary antibodies were: NONO (PAJ818Hu01, Cloud-Clone Corp., Houston, TX), HAND2 (E-AB-61488, Elabscience, China), MYCN, PHOX2B, β-actin (AF5204, DF9730, AF7018; Affinity Bioscience, China).

### Cytotoxicity assays

Five thousand cells/well were plated into 96-well plates (Biolot, Russia) and left overnight. Auranofin was tested at concentrations 0.1-50 µM. Cells were treated for 72 h at 37 °C, 5% CO_2_. Cell viability was determined by 3-(4,5-dimethylthiazol-2-yl)-2,5-diphenyltetrazolium bromide assays (MTT, Sigma-Aldrich, Burlington, MA). Absorbance was registered on a Tecan spectrophotometer (Tecan Spark™, Switzerland) at 570 nm. The percentages of viable cells were calculated as a ratio (OD_570_ in wells with respective auranofin concentrations) to (OD_570_ of vehicle-treated cells) x 100%.

For colony formation assays, cells were transfected with the respective siRNAs (indicated in Results) for 24 h, then treated with 0.25 uM auranofin for 24h plated into 60 mm Petri dishes (100 cells/well, 10 ml of culture media) and incubated at 37°C, 5% CO2 for 14 days. Colonies were fixed in methanol, stained with 0.05% crystal violet for 30 min and counted.

For cell cycle analysis, cells were treated as indicated in *Results*, then detached and collected by centrifugation. Pellets were washed twice with PBS, fixed with 70% ethanol in PBS at −20°C for 30 min, treated with 25 µg/mL RNAse A, stained with 20 µg/mL propidium iodide (PI) for 15-30 min, resuspended in PBS and analyzed on a CytoFlex B2-R2-V0 flow cytometer (Beckman Coulter, Brea, CA) at excitation/emission settings 544/570 nm (TRITC channel, 532 nm laser). The loss of plasma membrane integrity (necrotic cell death) was determined by the percentage of PI-positive cells after treatment with 0.5 μM auranofin for up to 24 h, washing with PBS and staining with 50 µg/mL PI at room temperature for 30 min followed by flow cytometry (TRITC channel). Twenty thousand fluorescence events were collected per each sample.

### Statistical analysis

The GraphPad Prism 8.0 program (GraphPad Software, Boston, MA) was used to plot graphs. Results were processed using a one-way ANOVA test. Differences were statistically significant at *P ≤ 0.05, **P ≤ 0.01. At least three biological replicates were used per each experiment.

## Results

### Auranofin and siNONO elevate intracellular ROS

Auranofin, an inhibitor of TrxR1, has been implicated in inactivation of NONO protein in neural cells [42, 45]. We hypothesized that this effect is associated not as much with direct NONO inhibition by the auranofin ligand but instead with the shift of intracellular redox balance in response to TrxR1 inhibition. To test this assumption, we examined the effects of auranofin on intracellular ROS generation using flow cytometry-assisted detection of DCFH_2_-DA dye fluorescence. H_2_O_2_ (0.5 mM) was used as a control for intracellular ROS accumulation [46]. As shown in Figures 1A and S1, auranofin (0.5 μM) rapidly increased the portion of DCF-bright cells in NB cells (Kelly, IMR-32 and SK-N-AS) and, to a lesser extent, in non-malignant HS5 fibroblasts. Importantly, treatment with siNONO phenocopied the ROS-inducing effect of auranofin. Moreover, the combination of 0.5 μM auranofin and siNONO further elevated (up to ~50-75%) the percentage of bright cells in Kelly and IMR-32 cell lines. In SK-N-AS and HS5 counterparts, this combination was less efficient. Intracellular oxidation was associated with a significantly decreased abundance of *TrxR1* and *NONO* mRNAs as early as by 2 h of cell exposure (Figure 1B). In contrast, the steady-state levels of *CDKN1A* mRNA increased ~3-4-fold in response to auranofin, arguing against general down-regulation of transcription in auranofin-treated cells. Together, these data implicated NONO in the control of intracellular redox status, a previously unmet function.

**Figure 1.**
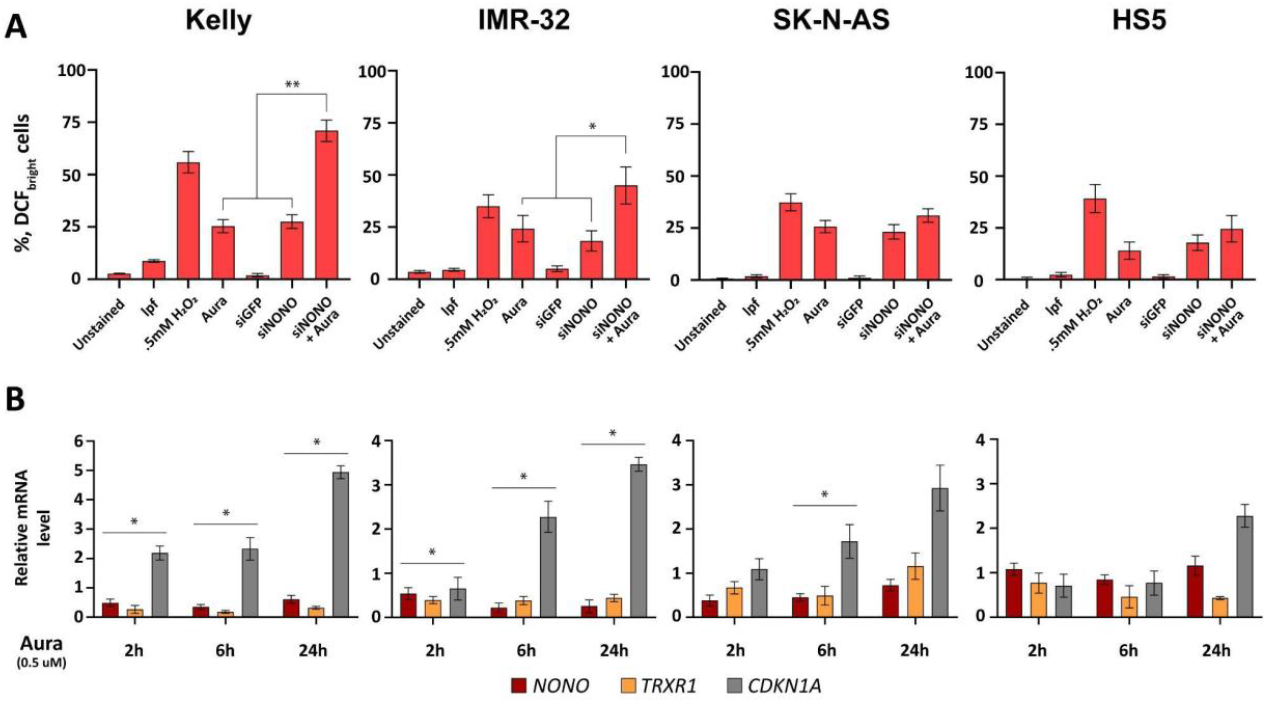
Effects of auranofin (Aura) and *NONO* knockdown on intracellular ROS content and gene expression. Cells pre-treated with DCFH_2_-DA were exposed to the indicated agents and analyzed by flow cytometry (see *Materials and Methods* for details). (A) Percentages of cells with elevated ROS. lpf, liposomes. (B) Time course of steady-state levels of *NONO, TRXR1* and *CDKN1A* (normalized to *HPRT* message) in response to 0.5 μM auranofin. Levels of each transcript in untreated cells were taken as 1 (control). Bars above the column groups show statistical significance relative to the control. Data are mean ± standard errors (n=3). *P ≤ 0.05, **P ≤ 0.01.

### Individual CRC are differentially regulated by auranofin and siNONO

Next, we were interested in (1) whether individual CRC genes are differentially regulated by auranofin-induced ROS elevation in three NB cell lines *vs*. non-malignant fibroblasts, and (2) is there a role of MYCN in this regulation? As shown in Figure 2A, treatment with gene-specific siRNAs was efficient: siMYCN attenuated *MYCN* messages ~2-3 fold; a similar extent of *NONO* down-regulation was achieved with siNONO. Importantly, auranofin and MYCN/NONO siRNAs differentially influenced the expression of *HAND2* and *PHOX2* genes. In Kelly and IMR-32 cells, auranofin activated these genes ~ twofold, and this activation was prevented by siNONO but not by siMYCN. In SK-N-AS and HS5 cells, auranofin evoked minor effects (<1.5 fold) on *HAND2* and *PHOXB* mRNAs; again, siNONO attenuated the steady-state levels of these transcripts (Figure 2A). Results of RT-PCR experiments were replicated at the protein level (Figure 2B). The analysis of cell cycle distribution showed that the *MYCN*-amplified Kelly was sensitive to siRNA-mediated *NONO* suppression: the portion of nuclei with fragmented DNA (subG1 events) increased up to 60%), in contrast to cell lines carrying single *MYCN* copy (Figures 2C and S2). Thus, auranofin and *NONO* knockdown synergized in the decrease of the factors relevant to NB cell survival. These results led us to test the therapeutic meaningfulness of these combinations.

**Figure 2.**
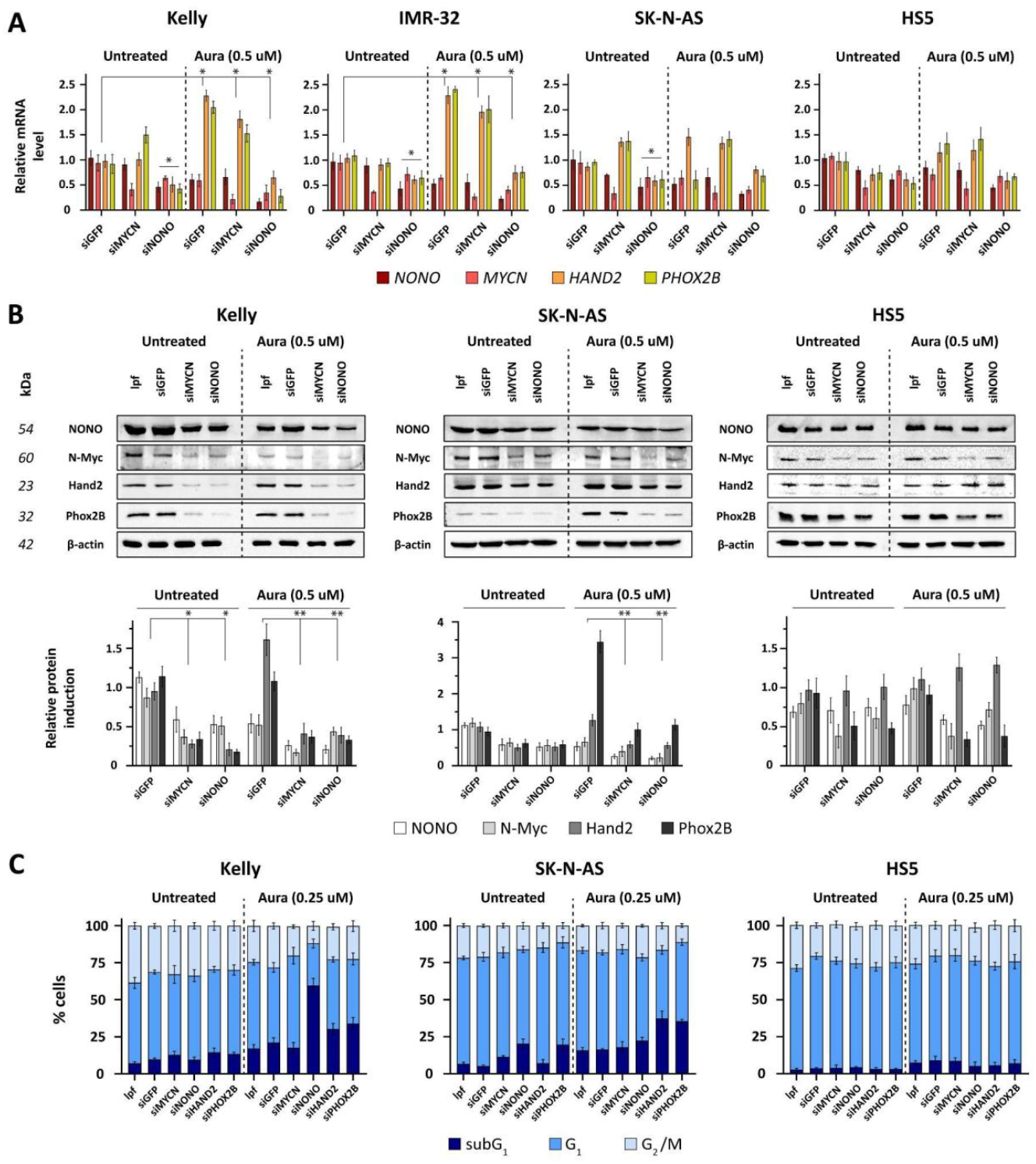
Auranofin, NONO and MYCN cooperate for CRC deregulation in NB cell lines. Steady-state levels of CRC mRNAs (A) and proteins (B). (A) Gene-specific signals were normalized to *HPRT* transcripts. The ratio in untreated cells was taken as 1 (control). Bars above the column groups show statistical significance relative to the control. (B) *Top*, immunoblotting of cell lines transfected with siMYCN or siNONO followed by exposure to auranofin. Loading control: β-actin. *Bottom*, densitometric analysis of blots. Levels in mock-transfected (lpf only) cells were taken as 1. *P ≤ 0.05, **P ≤ 0.01. (C) Cell cycle distribution after treatment with CRC siRNAs ± auranofin. Shown are mean values and standard errors (n=3). See *Materials and Methods* for details. Aura, auranofin; lpf, liposomes.

### Auranofin is preferentially cytotoxic for MYCN-amplified Kelly cells: synergy with NONO knockdown

We addressed the question whether intracellular oxidation evokes death of NB cells with different *MYCN* status. Figure 3A showed that auranofin was cytotoxic for NB cell lines; by 72 h of treatment the mean IC_50_ values varied from 0.39 µM for *MYCN*-amplified Kelly cells to 1.2 µM for SK-N-AS cells with single *MYCN* copy. The HS5 fibroblasts were the least sensitive (mean IC_50_ 1.9 µM). Differential sensitivity of NB cells was further demonstrated by the cell cycle analysis (Figures 3B and S3): auranofin elevated the percentage of subG1 events in a dose-dependent manner. By 24 h this effect was the biggest in *MYCN*-amplified Kelly cells: ~30-40% and 50-60% for 1 μM and 2 μM auranofin, respectively. In IMR-32 cells, subG1 portions were less pronounced. In SK-N-AS and HS5 cells, the respective percentages were only 10-20%. In the most sensitive Kelly cell line, DNA fragmentation was accompanied with moderate and relatively late loss of the plasma membrane integrity, with ~20% PI-positive cells by 24 h (Figure 3C), suggesting an apoptotic mode of cell death.

**Figure 3.**
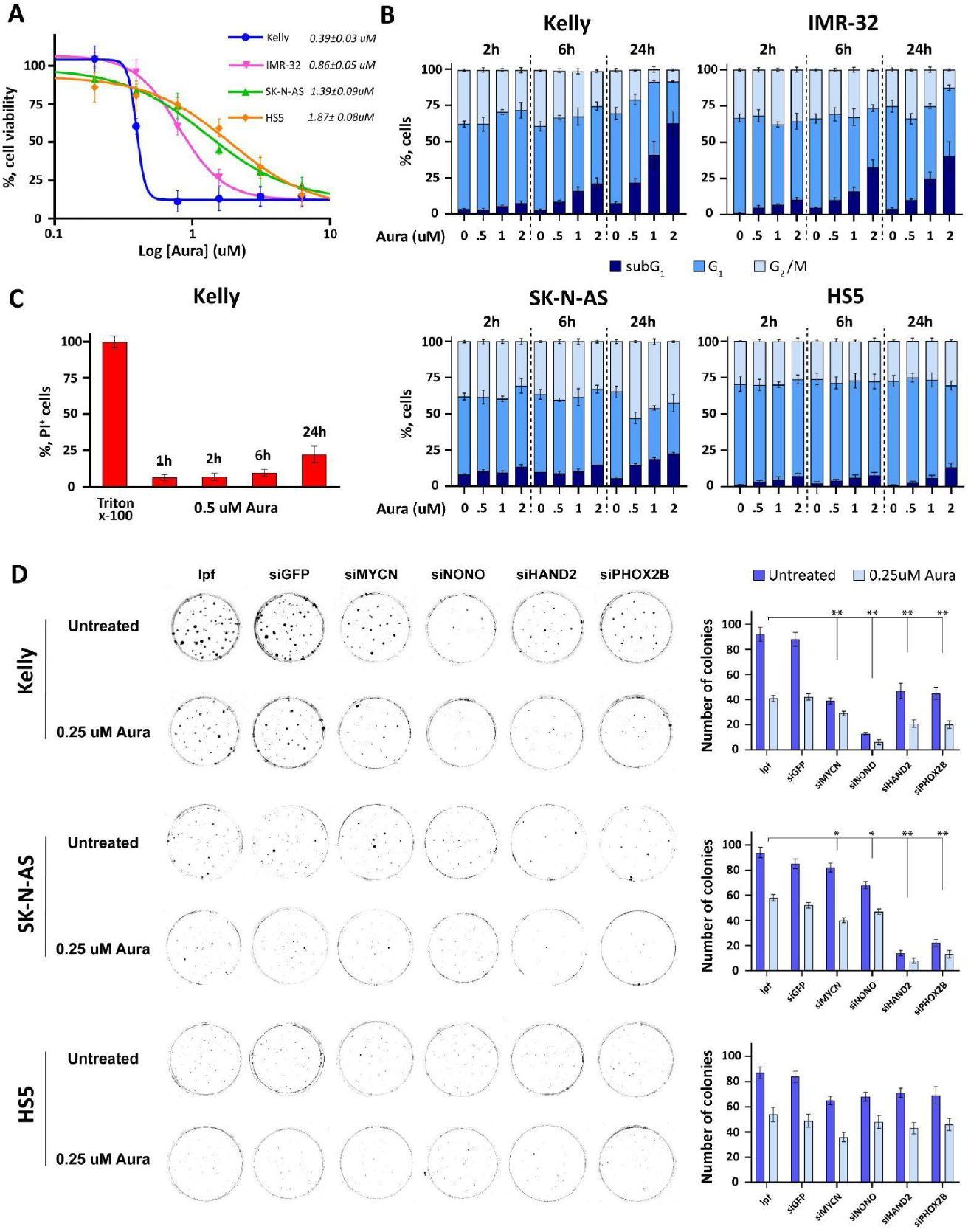
Cytotoxicity of auranofin combined with knockdown of individual CRC genes. (A) Dose response of NB cell lines and HS5 fibroblasts to auranofin (72 h, MTT assay). (B) Cell cycle distribution in auranofin-treated cells. Shown are mean ± standard errors, n=3, 20,000 events per sample. (C) PI positivity in Kelly cells. Triton X-100 (0.1%) was added as a positive control for PI entry. (D) Colony formation assays 14 days after transfection with the respective siRNAs and treatment with auranofin. Data are mean ± standard errors (n=3). See *Materials and Methods* for details.

Since auranofin and siNONO synergized in down-regulation of *MYCN, HAND2* and *PHOX2B* genes (see above), we were interested in whether these combinations increase NB cell death. Unlike shorter treatments, in these experiments the readout of the cytotoxic potency was single cell survival determined by colony formation 14 days post treatment. Figure 3D shows that, alone, siRNAs NONO, MYCN, HAND2 and PHOX2B decreased the clonogenic survival whereas the control siGFP did not. Most importantly, each knockdown was synergistic with sublethal (0.25 μM) concentration of auranofin. As in other assays, the most pronounced efficacy was observed in Kelly cells: siNONO was the most potent alone (~10-fold reduction of colony numbers *vs*. vehicle control) and together with auranofin (~20-fold reduction). In SK-N-AS cells, siRNAs against *HAND2* and *PHOX2B* showed the best inhibitory effects (5-7-fold decrease of colony numbers). In HS5 fibroblasts, the cytotoxicity of auranofin, alone or together with each CRC-specific siRNAs, was significantly lower compared to NB cell lines.

In summary, intracellular ROS generation by submicromolar concentrations of auranofin potently down-regulated the expression of the *MYCN, NONO, HAND2* and *PHOX2B* genes in NB cell lines, the *MYCN*-amplified Kelly cell line being the most sensitive. Although each of these genes is differentially regulated by auranofin, the combinations of this drug with knockdown of *NONO* (Kelly) or *HAND2* (SK-N-AS) substantially increased the cytotoxic potency. These data provide evidence in support of CRC members, in particular, NONO, as novel regulators of redox status in NB cells.

## Discussion

In this work we identify the multifunctional RNA/DNA-binding protein NONO as a previously unrecognized sensor of intracellular oxidation in NB cells. Transient *NONO* inactivation mimicked ROS generation in response to the TrxR1 antagonist auranofin. Combinations of *NONO* knockdown and auranofin were preferentially cytotoxic for *MYCN*-amplified (ADRN state-related) NB cells. Two lines of evidence support this conclusion: first, auranofin rapidly decreased the abundance of *NONO* mRNA and protein. Second, siNONO phenocopied ROS elevation by auranofin. Intracellular oxidation down-regulated CRC genes *HAND2* and *PHOX2B*. Most importantly, combinations of auranofin and siNONO potently reduced long-term survival of *MYCN*-amplified cells while evoking less pronounced effect on NB cells with single *MYCN* copy and sparing non-malignant counterparts. These observations provide evidence of a bidirectional link between NONO and cellular redox homeostasis: oxidation attenuates *NONO* mRNA / protein and, conversely, *NONO* knockdown exacerbates intracellular oxidation.

Mechanistically, several not mutually exclusive models may account for the observed phenomena: (i) direct modification of NONO (e.g., oxidation of redox-sensitive cysteine residues) could impair its binding to nucleic acids or promote proteasomal degradation; (ii) ROS-dependent limitation of availability of NONO-interacting partners (splicing factors, paraspeckle components, or RNA helicases) may reduce NONO stability or function; (iii) NONO loss may derepress transcriptional programs that activate mitochondrial respiration or diminish the antioxidant capacity (for instance, transcriptional down-regulation of glutathione peroxidase 1 in glioblastoma [47]), thereby producing a feed-forward ROS increase. Distinguishing between these possibilities will require direct assays (mass spectrometry mapping of post-translational oxidative modifications on NONO, redox-dependent co-immunoprecipitation, and measurements of NONO-RNA binding under oxidizing conditions) as well as genetic rescue experiments using oxidation-resistant NONO mutants.

It is known that *MYCN* amplification confers a transcriptional and metabolic state characterized by heightened anabolism and dependence on robust antioxidant systems [48].

Our observation that *MYCN*-amplified cells were more sensitive to auranofin and siNONO is consistent with the hypothesis that CRC integrity and redox status are interdependent vulnerabilities in the ADRN NB. NONO’s role in the maintenance of CRC circuit and association with *MYCN* mRNA [49] surmises that NONO suppression can weaken the transcription-driven proliferation and impair adaptive responses to oxidative stress. The clonogenic loss by auranofin-siNONO combination likely reflects a collapse of lineage-defining transcription and compensatory antioxidant defenses in cells metabolically stressed by *MYCN* amplification. These findings highlight NONO as an exploitable intersection of transcriptional tuning for adaptation to redox imbalance.

Auranofin has a clinical history as an anti-rheumatoid drug owing to its potent and irreversible inhibition of TrxR1 [50]. As a repurposing candidate, auranofin has been explored in cancer treatment. Our data support the concept that transient induction of oxidative stress can lead to targeted destabilization of individual CRC members (HAND2, PHOX2B) to achieve a pronounced long-term lethality in *MYCN*-amplified NB. Importantly, auranofin has been reported to inactivate NONO at least indirectly as a part of metabolic response to intracellular oxidation [45]. Recent discovery of (R)-SKBG-1 compound that covalently binds to a critical cysteine residue in NONO [35] provided a unique instrument for direct pharmacological NONO inactivation. It is of a particular interest to investigate whether (R)-SKBG-1 evokes similar effects on ROS content and CRC gene expression as auranofin and siNONO. Also, protein degraders using (R)-SKBG-1 structure for NONO recognition (PROTAC-like approaches) are likely to be feasible. The fact that auranofin is an approved drug could accelerate early clinical testing of combinations given that safety considerations— particularly systemic oxidative stress and off-target toxicities—are rigorously evaluated. Notably, our data suggest a certain selectivity, as non-malignant HS5 fibroblasts were less susceptible to the combination treatment.

NB displays lineage plasticity between ADRN and MES states, and treatment-induced selection or trans-differentiation toward MES phenotypes has been associated with resistance [15]. Oxidative stress, a condition that accompanies NB growth in a mouse model [51], can favor state transition or select for MES-like cells that survive ROS insult [52]. Alternatively, ROS burst may activate compensatory transcriptional programs that involve *HAND2/PHOX2B* before terminal collapse. Future work should address the question of whether the surviving cells acquire MES phenotype(s) and evaluate the regimens that impede plasticity by combining NONO attenuation with inhibitors of pathways that support MES survival. Likewise, up-regulation of alternative antioxidant systems (glutathione biosynthesis, NADPH-producing enzymes) may produce resistance; profiling adaptive transcriptomic and metabolomic responses will be essential to design durable combinations.

Our study is restricted to a limited set of cell lines and relies on transient *NONO* knockdown and pharmacological TrxR1 inhibition. These constraints prevent definitive conclusions about long-term consequences, *in vivo* tolerance, and effects on tumor microenvironment and immune components. Although the synergy in clonogenic assays was reproducible, the precise mechanism(s) whereby NONO attenuation elevates ROS remains to be elucidated. Finally, the possibility of adverse effects in ROS-sensitive tissues, such as neurons and heart, necessitates a cautious evaluation in animal studies.

In conclusion, we identify NONO as a mechanistic link between the redox status and CRC maintenance, particularly in *MYCN*-driven NB. This finding provides proof-of-principle for targeting CRC transcription by oxidative stress to produce marked cytotoxicity (Figure 4). Such a dual-strike strategy exploits an intrinsic vulnerability of transcriptionally compromised tumors and is attributable to other cancers with similar dependencies. Systematic dissection and rigorous preclinical testing are warranted to pave the way for repurposing of redox modulators along with the emergence of NONO-targeted therapeutic modalities in high-risk NB.

**Figure 4.**
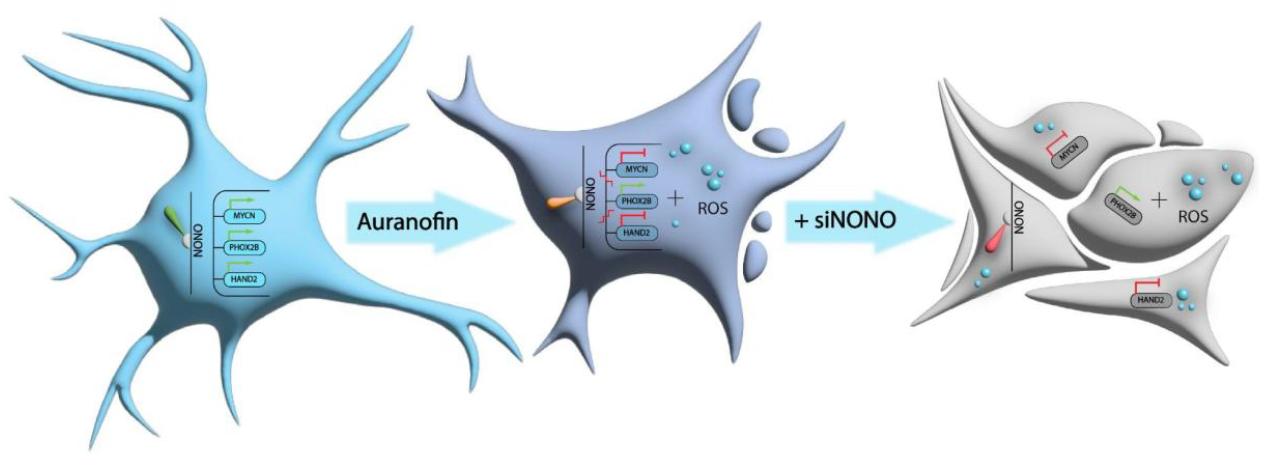
Auranofin and *NONO* knockdown synergize in triggering intracellular oxidation and death in NB cells. From left to right: intact neurocyte (blue); intracellular ROS generation and altered CRC gene expression in response to auranofin (pink); additional siNONO exposure leads to apoptosis as a result of redox imbalance (gray). Blue circles, ROS species; green lines, positive regulation; red lines, down-regulation.

## Abbreviations

ADRN: adrenergic
CRC: core regulatory circuit
GFP: green fluorescent protein
lpf: liposomes
NB: neuroblastoma
MES: mesenchymal
ROS: reactive oxygen species
siRNA: small interfering RNA
TrxR1: thioredoxin reductase 1.

## Funding

This study was supported by the Russian Science Foundation, grant 24-15-00097.

## Author Contributions

Conceptualization: A.A.S. and O.A.K. Methodology: S.S.P., O.O.M. and O.A.K. Validation: S.S.P. and O.O.M. Formal analysis: S.S.P., O.O.M. and O.A.K. Data curation: N.V.A. and A.A.S. Writing original draft: S.S.P., O.O.M., A.A.S. and O.A.K. Visualization: S.S.P. and O.A.K. Funding acquisition: N.V.A. Supervision: A.A.S. and O.A.K.

## Conflicts of Interest

The authors have no conflicts of interest to declare.

## Ethics Statement

Ethical approval is not applicable for this article.

## Notes

### Competing Interest Statement

The authors have declared no competing interest.

