## Supplementary material for "NONO as a Sensor of Intracellular Oxidation: Relevance to Neuroblastoma Cell Death": S1,

### Supplementary data

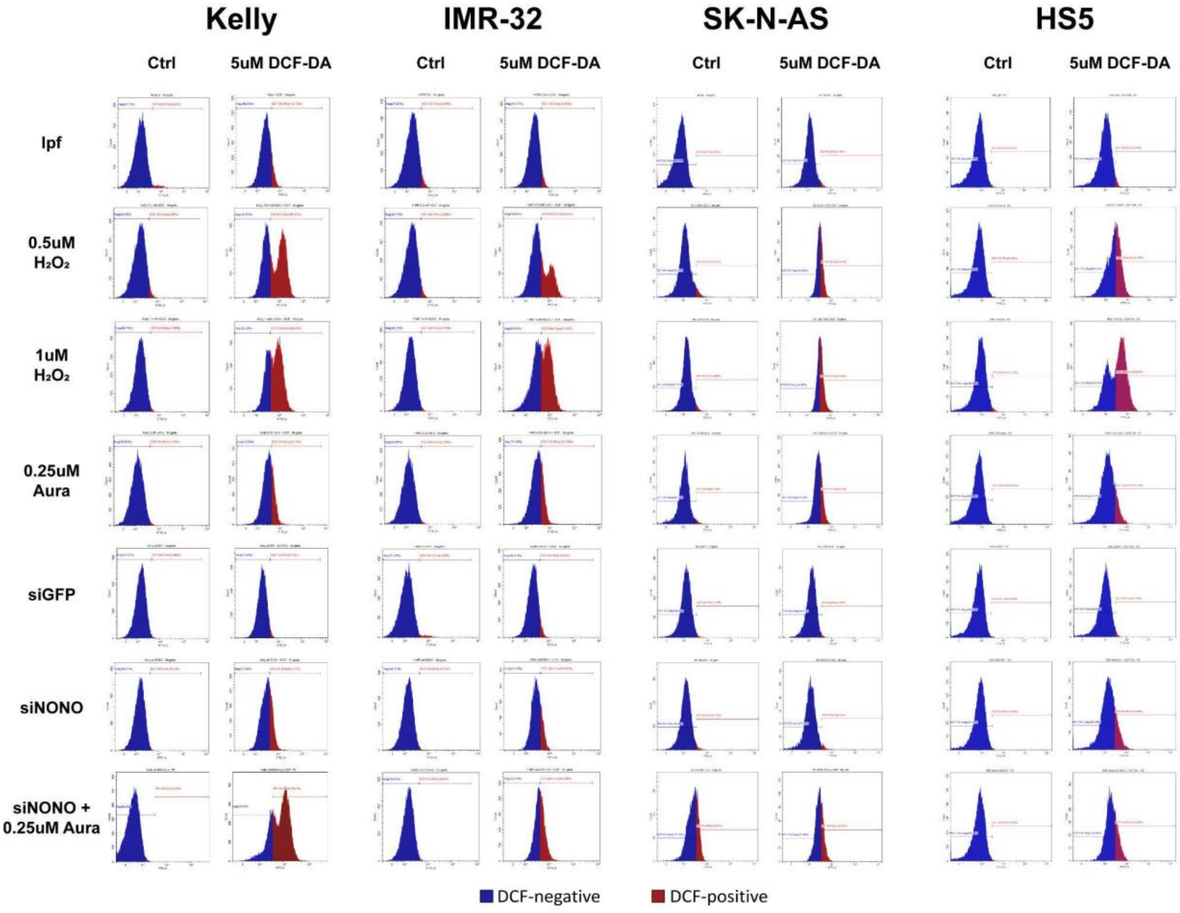


**Figure S1. Intracellular ROS content (DCFH$_2$-DA fluorescence assays).** See *Materials and Methods* for details.

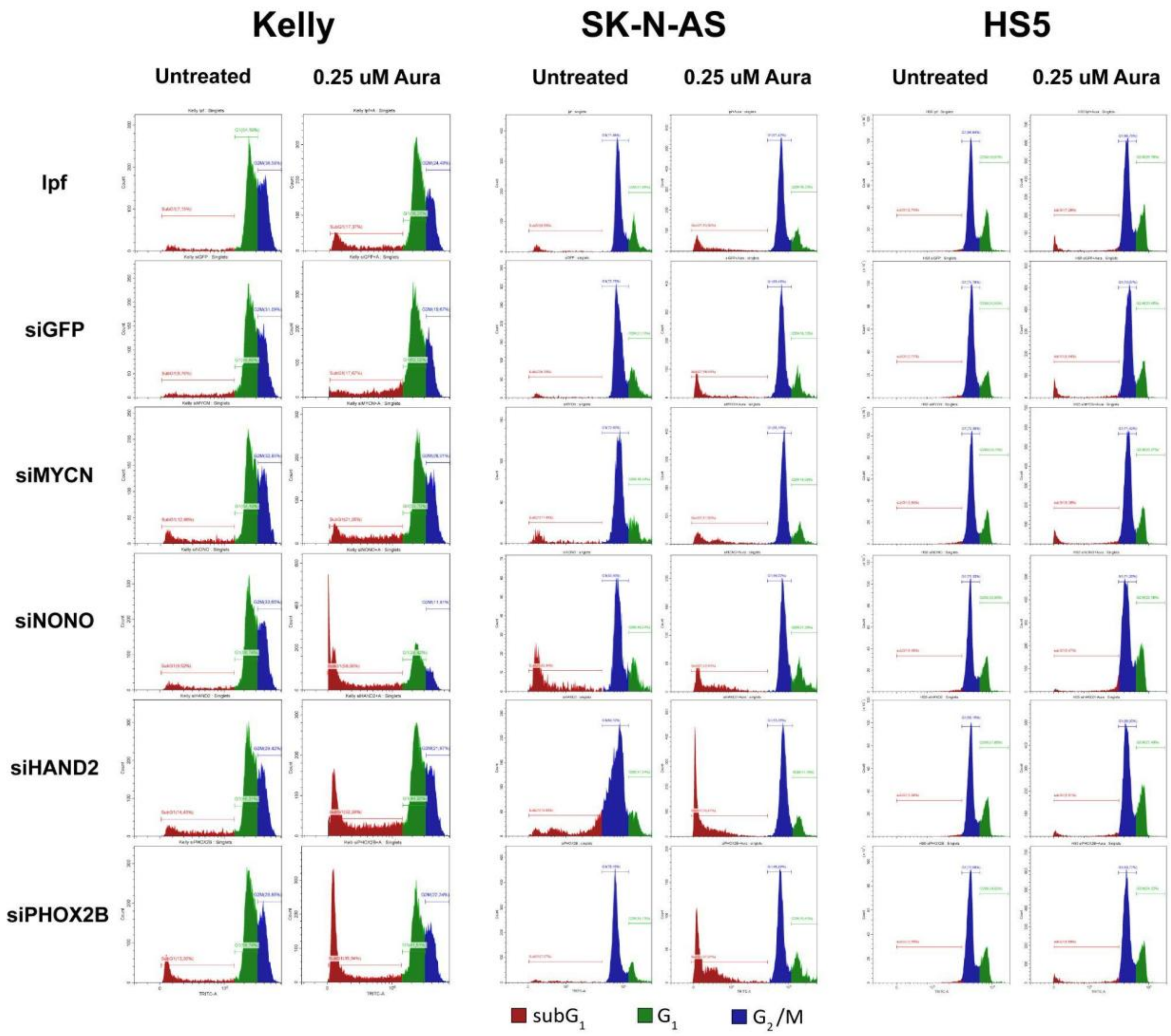


**Figure S2. Cell cycle distribution in response to combinations of individual gene knockdown and auranofin.** See *Materials and Methods* for details.

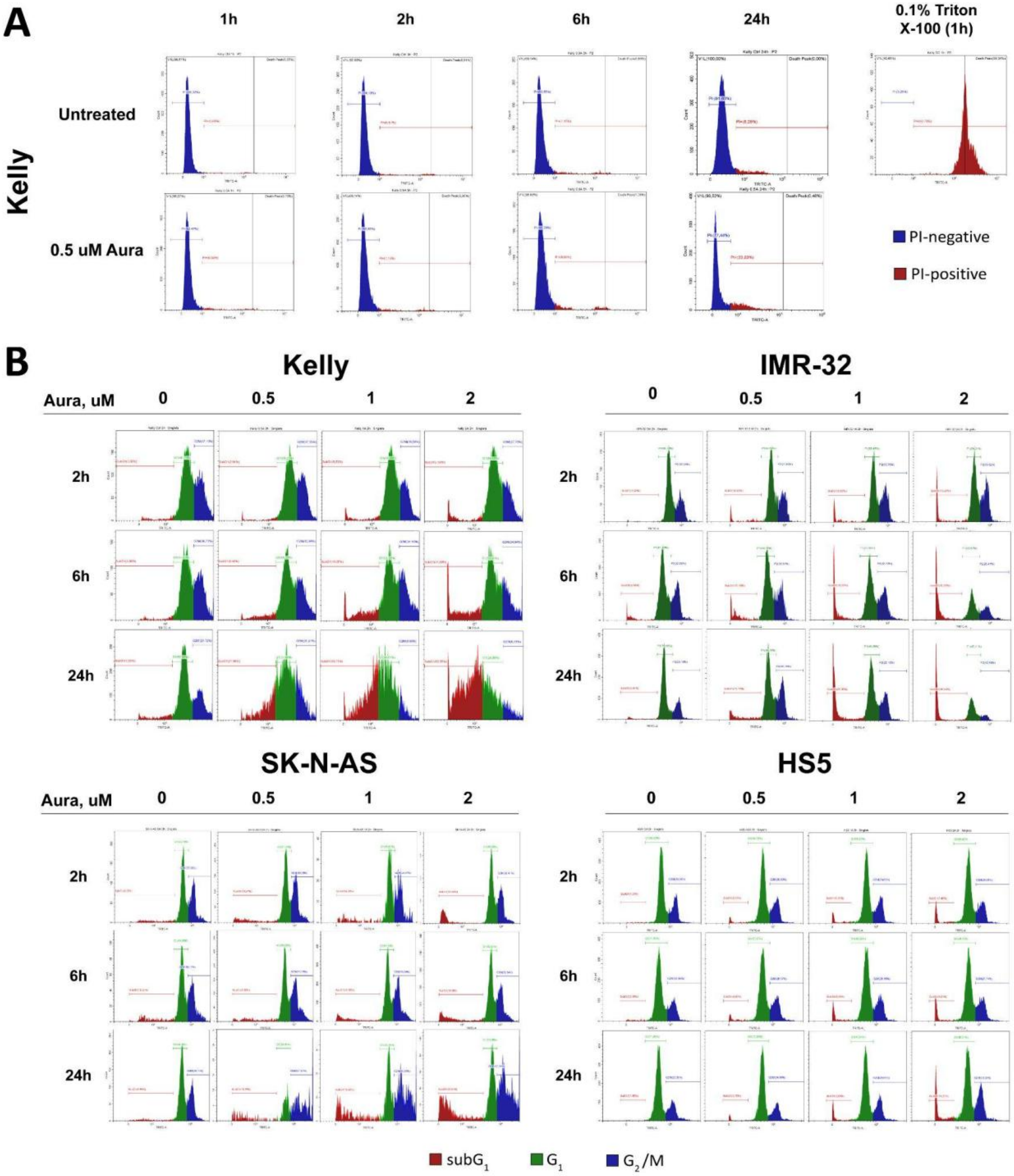


**Figure S3**. **Flow cytometry-assisted analysis of auranofin cytotoxicity.** (A) PI staining of whole cells; (B) cell cycle distribution. See *Materials and Methods* for details.
